# Glycation-lowering compounds inhibit ghrelin signaling to reduce food intake, lower insulin resistance, and extend lifespan

**DOI:** 10.1101/2022.08.10.503411

**Authors:** Lauren Wimer, Kiyomi R. Kaneshiro, Jessica Ramirez, Neelanjan Bose, Martin Valdearcos, Muniesh Muthaiyan Shanmugam, Dominique O. Farrera, Parminder Singh, Jennifer Beck, Durai Sellegounder, Lizbeth Enqriquez Najera, Simon Melov, Lisa Ellerby, Soo-Jin Cho, John C. Newman, Suneil Koliwad, James Galligan, Pankaj Kapahi

## Abstract

Non-enzymatic reactions in glycolysis lead to the accumulation of methylglyoxal (MGO), a reactive precursor to advanced glycation end-products (AGEs), which has been hypothesized to drive obesity, diabetes and aging-associated pathologies. A combination of nicotinamide, α-lipoic acid, thiamine, pyridoxamine, and piperine (Gly-Low) lowered deleterious effects of glycation by reducing MGO and MGO-derived AGE, MG-H1, in mice. Gly-Low supplementation in the diet reduced food consumption, decreased body weight, improved insulin sensitivity, and increased survival in leptin receptor-deficient (*Lepr^db^*) and wild-type C57B6/J mice. Transcriptional, protein, and functional analyses demonstrated that Gly-Low inhibited appetite-stimulating ghrelin signaling and enhanced the appetite-satiating mTOR pathways within the hypothalamus. Consistent with these molecular findings, Gly-Low inhibited ghrelin-mediated hunger responses. When administered as a late-life intervention, Gly-Low slowed hypothalamic aging signatures, improved glucose homeostasis and motor coordination, and increased lifespan, suggesting its potential benefits in ameliorating age-associated decline.

**Graphical Abstract:** 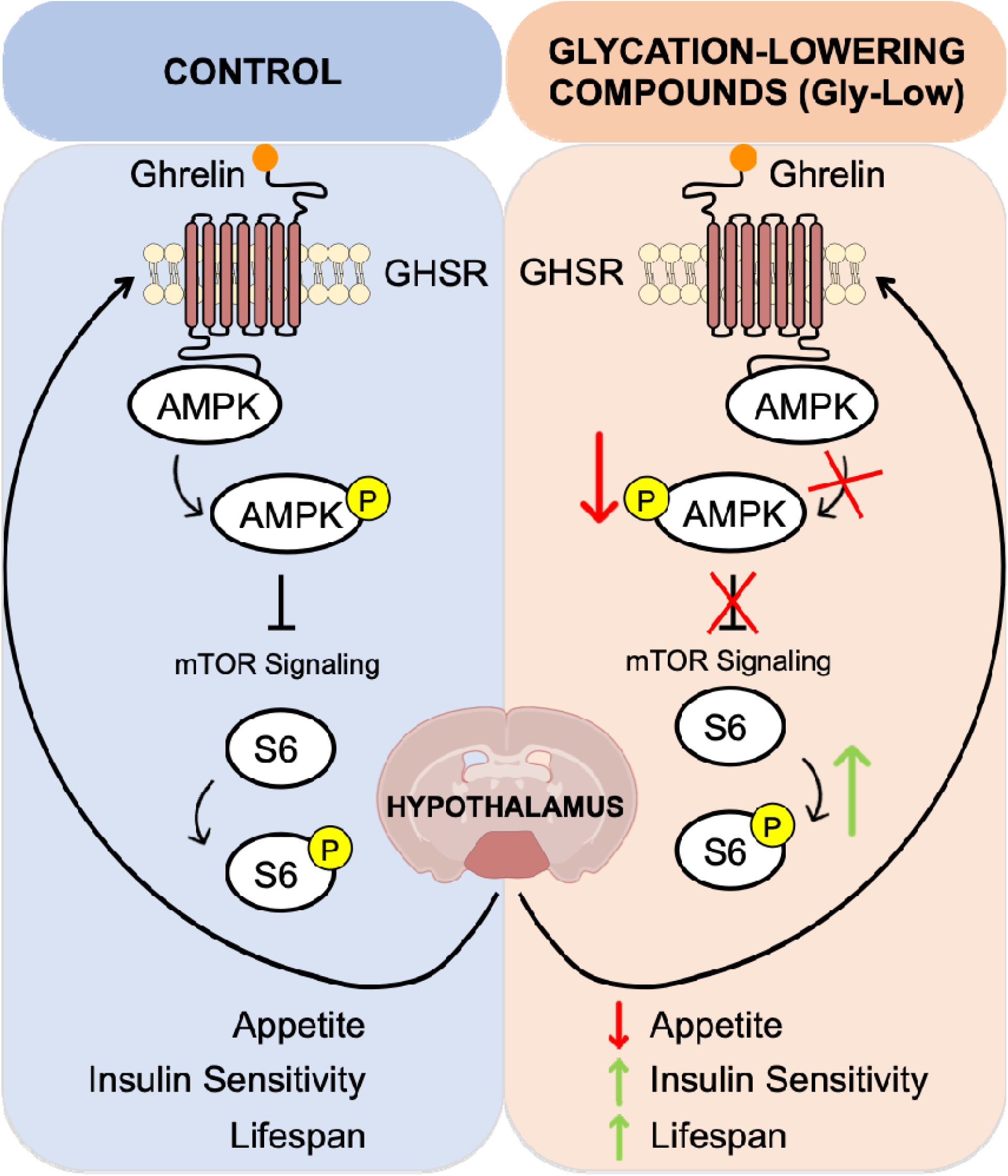

## Introduction

Despite the substantial efforts of public health, the incidence of obesity is growing worldwide^1^. Obesity reduces life expectancy by increasing the risk of several diseases^2^. Lifestyle changes have gained popularity to combat increasingly sedentary lifestyles and excess caloric intake, but dietary improvements remain challenging for most individuals^3,4^. Part of this challenge is due to homeostatic mechanisms governing food intake and energy expenditure to resist body weight loss^5^. Ghrelin and leptin are hormones that regulate food intake by promoting appetite and satiety, respectively^6–8^. Chronic dietary excess can disrupt leptin and ghrelin signaling impairing homeostatic regulation of food intake and thus promoting obesity^9^.

Food overconsumption and obesity are contributing factors to chronic hyperglycemia, which can alter glycolytic flux and thus increase the production of reactive α-dicarbonyls, such as methylglyoxal (MGO)^10,11^. MGO is an unavoidable byproduct of anaerobic glycolysis and is generated through the non-enzymatic degradation of glycolytic intermediates, dihydroxyacetone phosphate (DHAP) and glyceraldehyde-3-phosphate (G3P)^12^. Once formed, MGO reacts non-enzymatically with biomolecules such as proteins, lipids, and DNA to form advanced glycation end-products (AGEs)^13,14^. These covalent adducts have many deleterious consequences, such as impairing protein function, disrupting tissue architecture by forming extracellular crosslinks, activating inflammatory signaling cascades, altering cell-cell communication, and damaging nucleic acids^11^. Accumulation of these adducts, which occurs slowly with age or rapidly in hyperglycemic and obese individuals, drives many age-, obesity-, and hyperglycemia-associated pathologies. Cellular protection against AGEs occurs by endogenous glyoxalase enzymes, which detoxify MGO and thereby prevent downstream AGE formation^10,15^. We recently demonstrated using *C.elegans* that accumulation of AGEs also increases food intake^16^. We therefore hypothesized that therapeutically enhancing detoxification of AGEs may protect against obesity and obesity-associated pathologies.

### A Natural Compound Screen Identifies Compounds That Reduce Glycation Stress

Hyperglycemia, as often occurs in diabetes, accelerates AGE formation and its accumulation in various tissues, contributing to the development of diabetes-associated pathologies such as nephropathy, retinopathy, cardiomyopathy, neuropathy, and vascular injury^17^. Diabetic manifestations, such as peripheral neuropathy and reduced lifespan, were observed in the *C. elegans glod-4* mutant^18^, a model that lacks the endogenous detoxification pathway responsible for the clearance of MGO. We previously utilized this *glod-4* model to perform a high throughput screen of 640 natural compounds (TimTex, NPL640) to identify interventions to protect against AGE-associated pathologies and diabetic manifestations. We further tested 11 positive hits from our screen, as well as compounds from relevant glyoxalase-associated literature (thiamine^19^ and pyridoxamine^20^) in a cell culture model of glycation stress, as measured by rescuing neurite length retraction of rat dopaminergic (N27) neural cells following exposure to MGO (Figure S1A). We found that a combination of five compounds: α-lipoic acid, nicotinamide, piperine, pyridoxamine, and thiamine, termed Gly-Low, conferred better protection against MGO relative to the single compounds (Figure S1B).

Next, we tested Gly-Low in the context of obesity and hyperglycemia using a diabetic leptin receptor deficient (*Lepr^db^*) mouse model. These mice rapidly develop obesity, hyperglycemia, and glucose intolerance phenotypes that are accompanied by increases in glycation precursors and their adducts in several diabetes-relevant tissues such as the kidneys, heart, and liver^21^. Due to their early-onset diabetes, we started 8-week-old male *Lepr^db^* mice on a regular, low-fat diet supplemented with Gly-Low compounds for 16 weeks (Figure 1). LC/MS analysis of the plasma demonstrated significant reductions in absolute levels of both MGO (61% decrease) and its protein-bound arginine adduct, MG-H1 (41% decrease) in the plasma of mice fed a Gly-Low diet compared to controls (Figure 1A).

**Figure 1:**
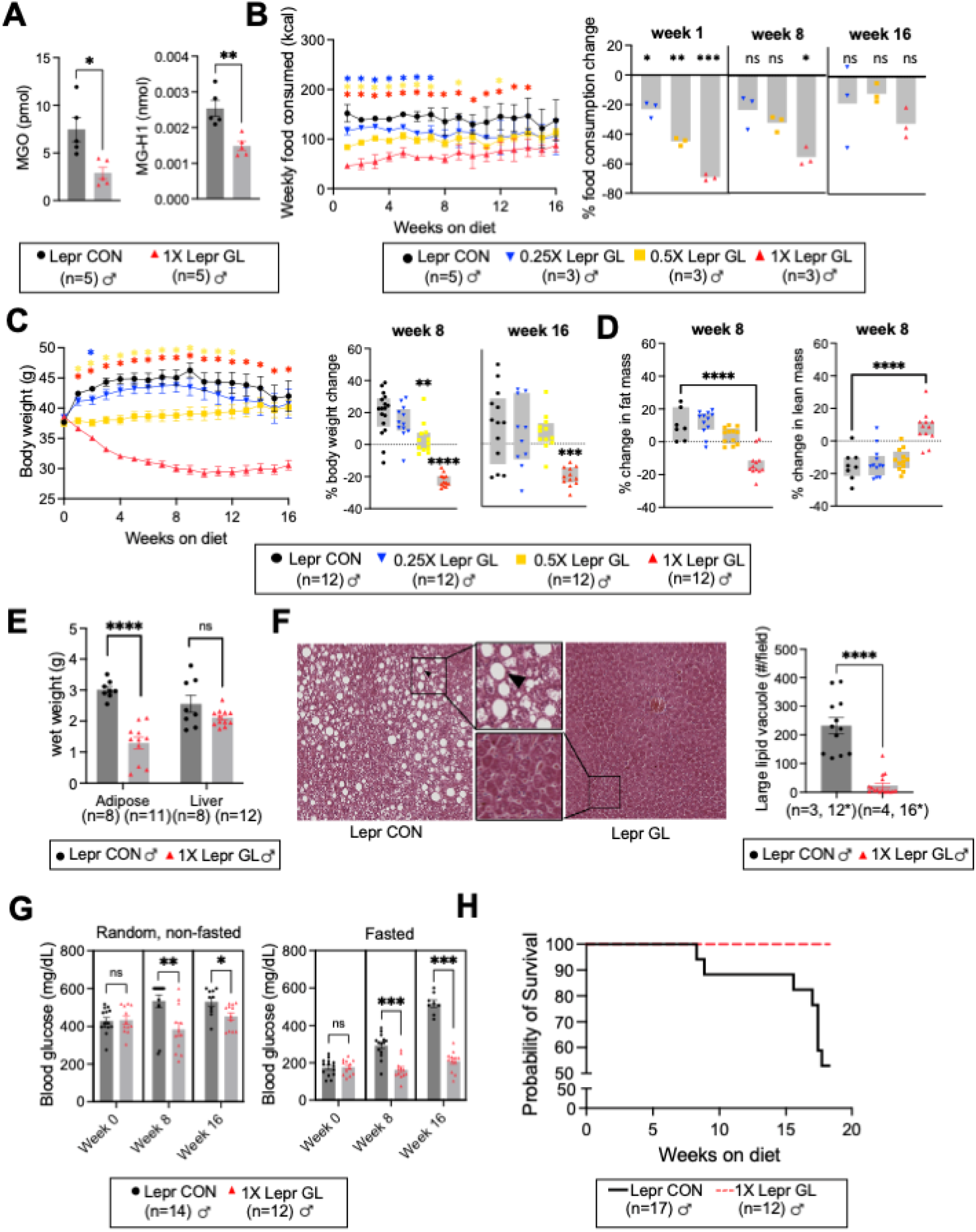
Glycation-lowering compounds (Gly-Low) lower glycolytic byproducts as well as rescue hyperphagia and obesity-associated pathologies in a diabetic, leptin receptor deficient (*Lepr^db^*) mouse model. A) Absolute levels of glycolytic byproduct, MGO (left), and its arginine adduct, MG-H1 (right), were reduced in the plasma of male mice treated with Gly-Low. n = 5 per treatment group, measured by LC/MS. Weekly food consumption (B) and body weights (C) of male *Lepr^db^* mice fed diets containing varying concentrations of Gly-Low were altered in a dosedependent manner. Food measurements of n = 3 cages of group-housed animals per treatment (n = 5 in the control group). Body weights of n = 12 mice per treatment group (n = 17 in the control group). Bar graphs (right) show food consumption differences (B) and percent change in body weight from starting diets (C) at their respective timepoint relative to control. D) Percent change from baseline in fat (left) and lean mass (right) of *Lepr^db^* mice on their respective diet. E) Wet weights of inguinal fat deposits and livers of *Lepr^db^* mice. F) Representative images of H&E stained liver sections from control and Gly-Low treated mice showing the presence of large lipid vacuoles (arrowhead). Quantification of liver lipid vacuoles, n = 3-4 livers per treatment group, 4* fields of view per animal. G) Random (nonfasted) and fasted (16h) blood glucose levels up to a max of 600 mg/dL. n = 12 per treatment group (n = 17 in the control, with deaths throughout the study [1 death before 8 weeks, 8 deaths before 16 weeks]). H) Survival curves of control mice and Gly-Low treated mice prior to their experimental endpoint. p = 0.007 (log-rank Mantel-Cox), p = 0.007 (Gehan-Breslow-Wilcoxon). Significance: ns (not significant), * p < 0.05, ** <0.005, *** <0.0005). Statistical analyses performed by unpaired T-test.

### Glycation-Lowering Compounds (Gly-Low) Reduce Food Intake and Diabetic Pathologies in *Lepr^db^* Mice

All concentrations of Gly-Low (0.25X, 0.5X, and 1X) reduced food consumption during the first 7 weeks on the diet, with the first 6 weeks showing a dose-dependent effect (Figure 1B). The failure to detect significant differences between the control group fed and those fed lower concentration Gly-Low diets (0.5X and 0.25X) in the latter half could be due to the increased variability in food consumption observed in the control group toward the latter half of the experimental timeline (Figure S2A). Gly-Low had an early dose-dependent effect on body weight gain that became insignificant for low-concentration diets (0.5X and 0.25X) as the experimental timeline progressed (Figure 1C). By 8 weeks of age, which is the age at which treatment began, *Lepr^db^* mice are already overweight. During the treatment period, control-fed *Lepr^db^* mice continued to gain body weight, reaching a group average of 45 grams by 8 weeks of treatment, roughly 50% more than an average wild-type mouse and representing a state of extreme obesity. In contrast, mice fed the highest concentration of Gly-Low diet lost an average of 22% of their starting body weight weights, reaching and maintaining a group average of 30 grams, which is roughly the average weight of wild-type (C57B6/J) male mice at that age^22^. Mice fed the 0.5X Gly-Low diet did not lose weight but did not gain weight over the experimental time course. The 0.25X Gly-Low diet did not prevent pathological weight gain of *Lepr^db^* mice.

Dual X-ray absorptiometry (DXA) is a direct measurement of bone and fat mass, with derived lean mass calculations. Using DXA methodology allowed us to determine whether the weight lost in Gly-Low treated mice was due to changes in the fat or lean mass compartments. After 8 weeks of treatment, control-fed mice gained an average of 10% of fat mass while 1X Gly-Low treated mice lost an average of 13% (Figure 1D, left). Additionally, the highest dose (1X) of Gly-Low was the only concentration sufficient to prevent the loss of lean mass (Figure 1D, right).

To assess whether changes in activity or metabolism could account for the weight loss observed in 1X Gly-Low treated mice, we performed metabolic cage testing and found no significant difference in activity or metabolic rate (Figure S2B and S2C). Given the pronounced effect of 1X Gly-Low treatment on reducing food intake, we performed several tests to rule out food aversion. When Gly-Low was administered via intraperitoneal (IP) injection, food consumption rates decreased similarly (Figure S3A-S3B). After an 18-hour fast, mice initially ate similar amounts of food whether given a low-fat Gly-Low or control diet (Figure S3C). When combined with a high-fat diet, a Gly-Low diet did not exhibite appetite-suppressive effects (Figure S3F), as seen in the low-fat diet (Figure S3E), suggesting that aversion is not due to palatability of the compounds. Conditioned taste aversion (CTA) was unexplored within our cohort, and the use of only male mice limits the assessment of sexual dimorphisms. These tests and the body weight results from long-term Gly-Low treatment (∼10 months) (Figure 2), suggest that Gly-Low was not taste aversive to mice. While Gly-Low had significant effects at several concentrations on both feeding rates and body weights, we restricted further *in vivo* analyses to the highest dosage of Gly-Low supplementation due to its effectiveness and robust changes in circulating AGE levels.

**Figure 2:**
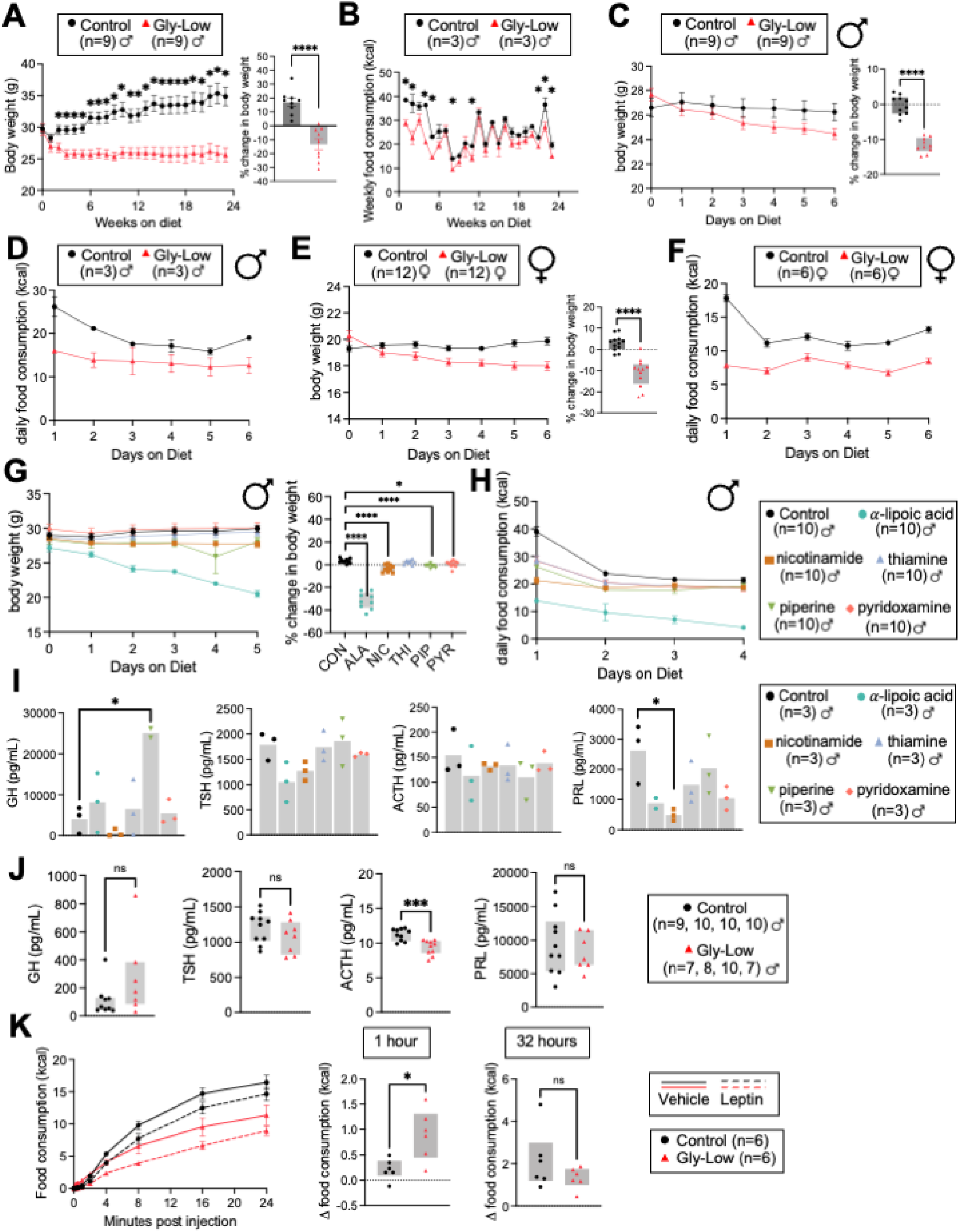
Gly-Low, and its constituents, reduce body weights and food consumption in male and female C57B6/J mice, independent of major pituitary hormones and hypothalamic leptin signaling. Body weights (A) and food consumption (B) of male wild-type mice chronically (24 weeks) treated with Gly-Low were reduced compared to control mice. Body weights (C) and food consumption (D) of male wild-type mice acutely (1 week) treated with Gly-Low were reduced compared to control mice. Body weights (E) and food consumption (F) of female wild-type mice acutely (1 week) treated with Gly-Low were reduced compared to control mice. Body weights (G) and food consumption (H) of male wild-type mice acutely (1 week) treated with individual compounds of Gly-Low compared to control mice. Graphs to the right demonstrate percent change in body weight from starting diets (A, C, E, G). Plasma levels of growth hormone (GH), thyroid stimulating hormone (TSH), adrenocorticotropic hormone (ACTH), and prolactin (PRL) from male mice acutely treated with Gly-Low constituents (I) and Gly-Low (J). K) Food consumption rates (kcal) of wild-type mice fed a control diet or a Gly-Low diet were reduced following an injection of leptin compared to those given a saline vehicle control. Bar graphs (right) show differences in food consumption between treatment groups at 1 hour and 32 hours after an injection of leptin compared to their saline food consumption. Significance: ns p > 0.05, * p < 0.05, ** <0.005, *** <0.0005). Statistical analyses performed by unpaired T-test.

Consistent with the reduced adiposity observed by DXA analysis, Gly-Low treated mice had significantly smaller fatty deposits in their inguinal fat pad (Figure 1E). The wet weights of dissected livers from Gly-Low treated mice were also significantly lower than those of control-fed mice (Figure 1E). Further histological assessment of livers from Gly-Low treated mice revealed an 89.8% reduction in large lipid vacuoles (Figure 1F). These changes in fat accumulation suggest that Gly-Low treatment improves lipid homeostasis observed in this mouse model.

In agreement with reduced glycation burden, Gly-Low treated mice displayed significantly improved glycemic control as indicated by a 9.9% reduction in fasted glucose and 28.4% reduction in non-fasted (random) blood glucose levels after 8 weeks of treatment that persisted over the entire course of treatment (Figure 1G). These observed improvements in glycemic control of Gly-Low treated mice were accompanied by reduced polyuria (Figure S2D), measured by weighing home cages to determine cage soiling. Polyuria occurs during diabetes when the kidneys fail to filter out sugar from the urine, resulting in excessive urine output^23^. This polyuria phenotype is largely driven by excessive blood sugar levels^23^. In mouse models of diabetes, excessive urine output can contribute to significant changes in cage weight due to excessive soiling of bedding and nesting material^24^. Thus, differences in cage weight can be used as a proxy for changes in polyuria. Water consumption rates of Gly-Low treated mice were measured during metabolic cage testing, and were not significantly different from rates of control-fed mice, indicating that reduced soiling was not due to reduced water intake (Figure S2E). In addition to polyuria, proteinuria, the presence of protein in the urine, indicates diabetic kidney damage^25^. We found that control mice had increasing protein levels in their urine over the experimental time course, which was indicative of worsening kidney function. In contrast, Gly-Low treated mice had reduced protein in their urine (Figure S2F), suggesting a protective role within the kidney.

The ability of Gly-Low to reduce hyperphagia and diabetes-associated pathologies in male *Lepr^db^* mice translated to a complete rescue in early mortality rates during the experimental timeline (Figure 1H). Previous reports demonstrated that male *Lepr^db^* mice had a median lifespan of 349 days, with mortality continuously occurring after 16 weeks of age. Consistent with these reports, we observed an 18% decrease in cohort numbers in our control-fed mice by 15 weeks on diet (23 weeks of age)^26^. By the time of dissections (26 weeks of age), control-fed mice had a 52.9% chance of survival, while Gly-Low treated mice had no recorded experimental deaths. Collectively, these findings in a leptin receptor-deficient mouse model highlight the therapeutic potential of Gly-Low in treating various obesity- and diabetes-associated conditions by reducing overconsumption, associated glycation burden, and glycation-associated pathologies.

### Gly-Low Reduces Food Intake In a Leptin-Independent Manner

Following experimentation in a leptin receptor deficient mouse model (*Lepr^db^*), we tested the highest dose of Gly-Low in a wild-type (C57BL/6J) mouse model. Long-term treatment of male wild-type mice demonstrated similar reductions and subsequent maintenance in body weights (Figure 2A) and food consumption rates (Figure 2B). Throughout the 6-month treatment, Gly-Low treated mice lost an average of 13.2% of their body weight, while control-fed mice gained an average of 16.9%. Similar to findings in the *Lepr^db^* mouse model, we used metabolic cage testing to assess mouse activity and metabolism and found that Gly-Low treated mice were losing weight primarily due to reduced food consumption rates and not due to changes in activity or energy expenditure (Figure S3G and S3H). To test for sex specificity, we tested the highest dose of Gly-Low as an acute (1 week) treatment and observed that age-matched, wild-type males and females demonstrated similar reductions in total body weights and food consumption (Figure 2C-2F). Young (3-month-old) males lost an average of 11.7% of their body weights, and young (3-month-old) females lost an average of 11.1%, indicating that the effect of Gly-Low on food consumption and body weight gain are not sexually dimorphic.

Next we determined which Gly-Low constituent was responsible for the observed effects. To test this, each compound comprising Gly-Low was incorporated into its low-fat diet at the same dose as in its final Gly-Low diet. Young (6-month-old) wild-type male mice were acutely treated with diets incorporated with α-lipoic acid (3 g/kg), nicotinamide (8.57 g/kg), thiamine (0.60 g/kg), piperine (0.26 g/kg), or pyridoxamine (2.43 g/kg). Interestingly, at the end of the single week of treatment, mice from each single-component group, except for thiamine, weighed significantly less than control mice (Figure 2G). Notably, mice treated with α-lipoic acid showed the most drastic effects on food consumption and body weights, consuming roughly 32% less food than control-fed mice (Figure 2H). This weight loss far exceeds that observed when α-lipoic acid is incorporated into the Gly-Low diet and may suggest a mechanism by which the other Gly-Low components dampen the effects of α-lipoic acid on appetite suppression and body weight loss. These findings indicate that the appetite suppression and weight loss observed in Gly-Low treated mice is largely driven by α-lipoic acid with weaker contributions due to nicotinamide, piperine, and pyridoxamine.

To address how Gly-Low components singly and in combination (in Gly-Low) may affect feeding behavior and body weight, we measured circulating hormone levels of the hypothalamic-pituitary axis known to have direct effects on body weight maintenance or feeding by performing a hormone multiplexing panel of growth hormone (GH), thyroid stimulating hormone (TSH), adrenocorticotropic hormone (ACTH) and prolactin (PRL). Specifically, during feeding, there is a reduction in GH secretion^27^. GH levels were significantly elevated in mice treated with piperine, while nicotinamide trended towards reducing its levels (Figure 2I, left). GH levels were unaffected when administered as a cocktail in Gly-Low (Figure 2J, left). These findings suggest that GH elevations induced by piperine are tempered by co-administration with the other components of Gly-Low. Hypothyroidism is often observed during reduced TSH production and release, leading to decreased basal energy expenditure and increased body weights^28^. Surprisingly, TSH levels had trending reductions in nicotinamide and α-lipoic acid treated mice (Figure 2I, middle left) and had no effect on TSH levels when administered in the Gly-Low cocktail (Figure 2J middle left). ACTH is a hormone produced and released by the pituitary gland responsible for regulating cortisol and androgen production^29^. Following feeding, ACTH is significantly reduced in circulating plasma levels^30^. We found that all of Gly-Low’s constituents trend towards reducing ACTH levels and, when given in combination (Gly-Low), caused a significant reduction in circulating levels of ACTH (Figure 2I and 2J middle right panels). These findings suggest additive effects of the individual constituents when co-administered in Gly-Low. We found that prolactin (PRL) levels are significantly lower in nicotinamide treated mice and trend lower in α-lipoic acid, pyridoxamine, and thiamine treated mice. However, PRL levels were unaffected when mice were treated with the cocktail of all constituents in Gly-Low. Collectively, our findings suggest that individual compounds of Gly-Low, when singly administered, differentially impact hormonal pathways that regulate feeding and body weight maintenance. Conversely, when these components are co-administered, the cockatil may impact these same pathways to a different degree.

Our collective findings assessing the individual components of Gly-Low suggest that nicotinamide, α-lipoic acid, piperine and pyridoxamine have significant effects on body weights. While appetite-suppression is largely driven by α-lipoic acid, our data suggests that other components, primarily nicotinamide, may drive body-weight effects independent of food consumption. The observation that after one week of treatment, nicotinamide-treated mice weighed less than control-fed mice is mirrored in our findings that *Lepr^db^* mice treated for 16 weeks with nicotinamide weighed significantly less than *Lepr^db^* mice fed a control diet (Figure S2G). This weight loss was not associated with signifianct changes in food consumption and indicates that in *Lepr^db^*mice, nicotinamide induces weight loss largely through mechanisms independent of food consumption and may suggest that a similar mechanism drives body weight changes in wild-type mice treated with nicotinamide.

Next, we investigated whether Gly-Low was also working indepependetly of the leptin signaling pathway in wild-type mice. To assess this, we subjected young (6-month-old), male wild-type mice to an injection of exogenous leptin and compared food consumption to saline-injected controls. Exogenous leptin stimulates pSTAT3 signaling within hypothalamic neurons, and a readout of this signaling can be measured by a reduction in feeding behavior. Similar to the control mice, Gly-Low treated mice demonstrated a decrease in food consumption when injected with leptin compared to saline injected controls (Figure 2K). Gly-Low treated mice appeared to have an increased difference in food consumption relative to control mice within 1 hour of the leptin injection, but significant changes in food consumption were lost by 32 hours post-injection. These findings indicate that leptin signaling is intact in Gly-Low treated mice and is further supported by a similar degree of pSTAT3 positivity in neurons within the arcuate nucleus of the hypothalamus in response to leptin injections (Figure S3I).

### Gly-Low Reduces Glycolysis and Enhances Cellular Detoxification Pathways in the Hypothalamus

The hypothalamus is aptly positioned at the brain-body interface (third ventricle) to sense circulating nutrients and hormones in the periphery to maintain body-weight homeostasis^31,32^. Thus, we chose to interrogate the hypothalamus for changes induced by Gly-Low molecularly. We assessed changes in hypothalamic transcripts using bulk RNA-sequencing and performed protein analysis on hypothalamic lysates from wild-type acutely (1 week) treated and control mice.

We investigated how glycation-lowering compounds influenced the expression of pathways relevant to glycation production and clearance. We found that Gly-Low treated mice showed a general reduction in the expression of glycolytic genes except for *Tpi* and *Pgk1*, whose increased activity is predicted to lower MGO production^33,34^ (Figure 3A and S4A). In contrast, genes directly responsible for MGO detoxification (glyoxalase genes) and a larger set of general cellular detoxification genes showed increased expression in Gly-Low treated mice (Figure 3A and S4A). Consistent with our transcriptome findings, targeted metabolomics data of plasma revealed decreased glycolysis and increased potential for MGO-specific and generalized cellular detoxification^35,36^ via elevated pentose phosphate pathway (PPP) utilization. The PPP regenerates NADPH, which is necessary for replenishing glutathione (Figure S4B). These findings suggest that Gly-Low may reduce the glycation burden by reducing its production and enhancing its clearance. Furthermore, these findings in the brains (hypothalamus) of wild-type mice complement our findings that Gly-Low reduces the systemic burden of the glycation precursor, MGO, and its predominant AGE, MGH1, in the blood of *Lepr^db^* mice.

**Figure 3:**
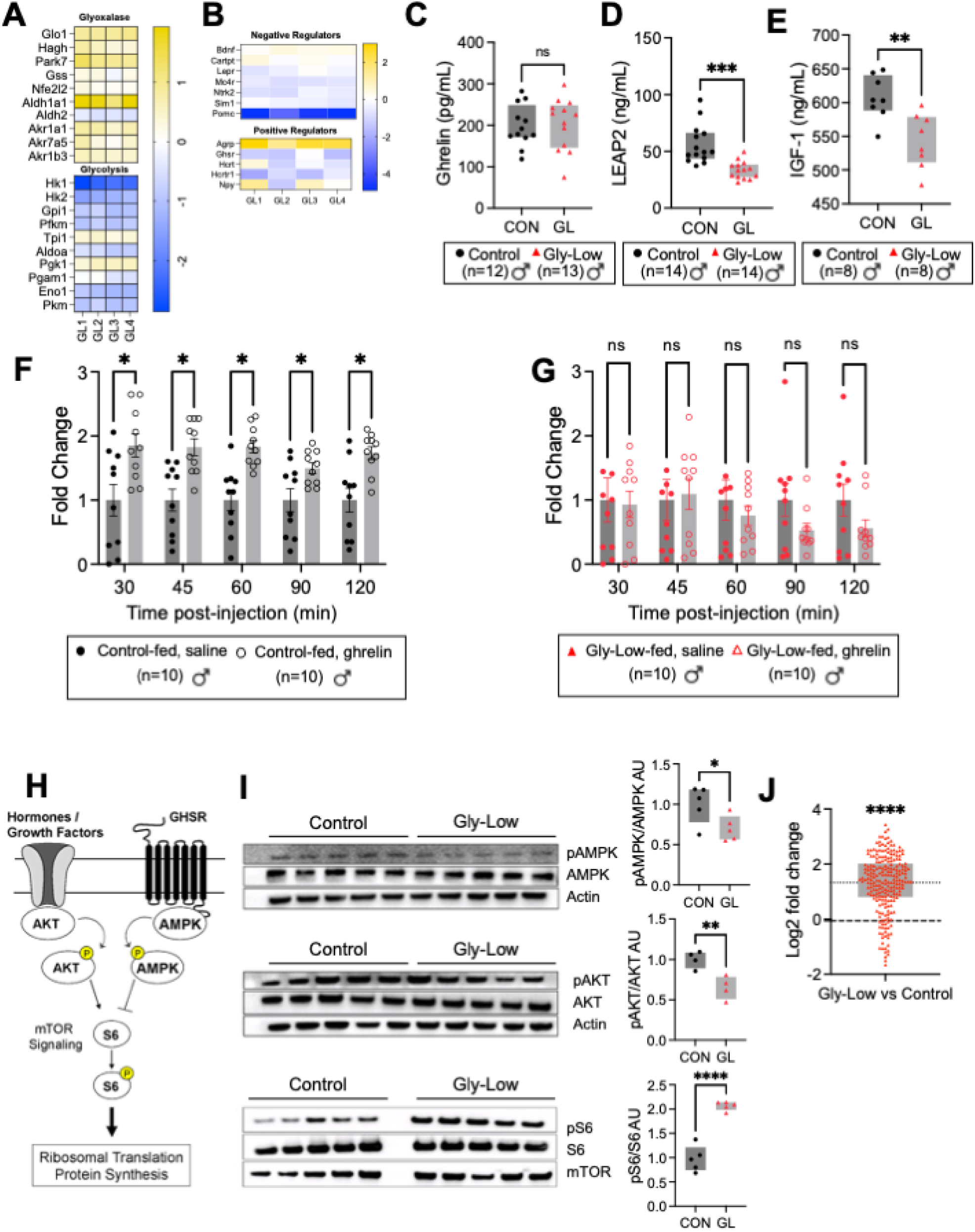
Gly-Low treatment alters hypothalamic genes responsible for appetite stimulation and blunts the efficacy of exogenous ghrelin signaling. A) Heatmaps showing transcript fold changes of genes involved in MGO production (glycolysis) and clearance (glyoxalase) in the hypothalamus of wild-type male mice acutely (1 week) treated with Gly-Low. B) Heatmap showing transcript fold changes of negative and positive appetite regulators in the hypothalamus of mice acutely (1 week) treated with Gly-Low. C) Plasma levels of acylated ghrelin were unchanged with Gly-Low treatment. Plasma levels of ghrelinreceptor antagonist, LEAP-2 (D), and ghrelin-stimulated growth factor, IGF-1 (E) were reduced with Gly-Low treatment. Exogenous ghrelin increased food consumption compared to saline injected controls in control (F) but not Gly-Low treated (G) mice. G) Schematic depicting activation or inhibition of mTOR signaling and ribosomal translation by hormone/growth factor or ghrelin signaling, respectively. H) Western blots of hypothalamic lysates from mice acutely (1 week) treated with Gly-Low. Protein expression of phosphorylated AMPK (top), phosphorylated AKT (middle), and phosphorylated S6 kinase (bottom) relative to their un-phosphorylated protein levels with quantification (right). I) Transcripts from ribosomal genes were significantly upregulated in hypothalamic lysates of mice acutely (1 week) treated with Gly-Low compared to control mice. Significance: ns p > 0.05, * p < 0.05, ** <0.005, *** <0.0005). Statistical analyses performed by unpaired T-test.

To investigate how Gly-Low may impact feeding behavior, we analyzed the 1,407 genes differentially expressed in the hypothalamus of Gly-Low treated vs control mice for changes to canonical feeding behavior genes. Interestingly, two well-characterized regulators of hunger and satiety, *Agrp* and *Pomc*, were consistently changed in the direction typically observed in hungry mice^37^ (Figure 3B). Analysis of this set of positive and negative regulator feeding genes in Gly-Low treated mice indicates that no one particular pathway was expressed at higher or lower levels than expected (Figure S4C).

### Gly-Low inhibits Ghrelin Signaling

The stomach-derived hormone ghrelin is a common signaling molecule to both the Agrp and the Pomc pathways. Ghrelin is a peptide produced in the stomach during fasting and times of hunger^38,39^. Following production, the circulating hormone binds its receptor, the growth hormone secretagogue receptor (Ghsr)^40^, within the hypothalamus to engage signaling pathways that stimulate food intake^41,42^. Ghrelin signaling is inhibited by competitive binding of the ghrelin receptor by the liver-derived hormone LEAP2^43^, or when ghrelin production is reduced during feeding^44,45^. To determine whether Gly-Low affected ghrelin production, we measured levels of acylated ghrelin and its endogenous antagonist, LEAP2, in the plasma. We found no significant difference in ghrelin levels between the treatment groups (Figure 3C). However, LEAP-2 levels were significantly lower in Gly-Low treated mice (40% lower) (Figure 3D), suggesting the absence of competitive binding of the Ghsr. Ghrelin signaling stimulates the release of growth hormone (GH)^39^, which in turn goes on to stimulate the release of IGF-1 in a well-studied GH/IGF-1 endocrine axis within the hypothalamus and pituitary^46^. To test if this ghrelin-related pathway is disrupted in Gly-Low treated animals, we measured insulin-like growth factor (IGF-1) in the plasma of control and Gly-Low treated mice. Consistent with disrupted ghrelin signaling, IGF-1 levels were significantly lower in Gly-Low treated mice relative to control mice (10.5% lower) (Figure 3E). To test if ghrelin signaling is indeed disrupted in Gly-Low treated animals, we subjected control and Gly-Low treated mice to exogenous acylated ghrelin, known to activate ghrelin signaling and increase appetite and induce feeding behavior^47^. In control mice, ghrelin-injected mice consumed significantly more food (average of 75% increase) relative to PBS-injected controls post-IP injection (Figure 3F). In contrast, Gly-Low treated mice failed to respond to ghrelin injections with increased food consumption (an average of 23% less food consumed relative to PBS-injected controls) (Figure 3G). These findings suggest a role for impaired ghrelin signaling in the reduced food intake observed upon Gly-Low treatment.

### Gly-Low Impairs Ghrelin Signaling and Increases mTOR Signaling In The Hypothalamus

Given that Gly-Low inhibits hypothalamic ghrelin signaling independent of ghrelin production, we investigated the biochemical changes downstream of the ghrelin receptor (Ghsr) (Figure 3H). Activation of the ghrelin receptor, Ghsr, as occurs during fasting, results in the increased phosphorylation of AMPK and downstream signaling pathways that stimulate appetite^48^. Chronic activation of AMPK has been shown to induce obesity and compromise beta cell function^49^. Growth factors, such as growth hormone and IGF-1, bind their respective receptors in the hypothalamus to counteract ghrelin by promoting phosphorylation of AKT. The phosphorylation of AKT promotes mTOR and subsequent S6 kinase phosphorylation and protein synthesis all of which are associated with nutrient signaling and appetite suppression^50^. Importantly, activated (phosphorylated) AMPK can directly and indirectly inhibit mTOR activity, highlighting the antagonistic relationship between these two pathways^51–53^.

Consistent with impairments in ghrelin signaling, we found that phosphorylated AMPK protein levels were significantly lower in hypothalamic lysates from Gly-Low treated mice and that levels of phosphorylated S6 protein were increased (Figure 3I). To further assess the activation of the mTOR/S6 signaling cascade, we assessed ribosomal transcripts from our hypothalamic RNA-seq dataset and found that nearly all ribosomal transcripts are upregulated (Figure 3J). This suggests that ribosomal biogenesis and, thus, protein translation, which occurs in response to mTOR/S6 signaling, is upregulated in the hypothalamus of Gly-Low treated mice. Further supporting this are the observations that *Fcf1* and *Nip7*, genes that encode proteins involved in ribosomal rRNA processing and critical to ribosomal biogenesis, were also significantly upregulated in our RNA-seq data set from Gly-Low treated mice (Figure S4D). Taken together, increased levels of phosphorylated S6 protein and transcript data to indicate downstream activation of ribosomal biogenesis are consistent with mTOR activation in the hypothalamus in response to Gly-Low treatment. Furthermore, phosphorylated AKT levels were significantly reduced in the hypothalami of Gly-Low treated mice (Figure 3I). This observation is consistent with our findings that IGF-1 levels, which can drive AKT phosphorylation, are also down in Gly-Low treated mice. Thus our observations that phosphorylated S6 protein levels are increased despite lower levels of phosphorylated AKT suggests that in the hypothalamus of Gly-Low treated mice, mTOR activity is increased in response to reduced inhibition by AMPK rather than stimulated by activation of AKT. Together, these experiments suggest that Gly-Low reduces food consumption by impairing appetite stimulating ghrelin signaling and activating appetite suppressive mTOR signaling^54,55^.

### Hypothalamic Transcripts of Gly-Low Treated Mice Do Not Mimic Those Of Caloric Restriction

Our studies identify that Gly-Low mimics traditional (enforced) caloric restriction (CR) phenotypes. While some of the benefits of Gly-Low can be attributed to the reduced caloric intake, leading to improved glucose homeostasis and reduced fat accumulation, it is worth noting the molecular differences between Gly-Low and CR. In comparing differentially expressed gene sets from the hypothalamus of Gly-Low treated mice with those from publicly available bulk-sequenced hypothalamus of calorically restricted mice^56^, we find negative correlations (p < 0.0001, R^2^ = 0.3923, Pearson’s correlation) in regressed fold changes (Figure S4E). Additionally, RNA sequencing of hypothalamic lysates of Gly-Low treated mice demonstrates a robust increase in ribosomal gene transcripts compared to control mice (Figure 4J), which is not observed in traditional CR. Thus traditional CR creates a state of hunger in mice while Gly-Low treated mice display signatures of satiated animals.

**Figure 4:**
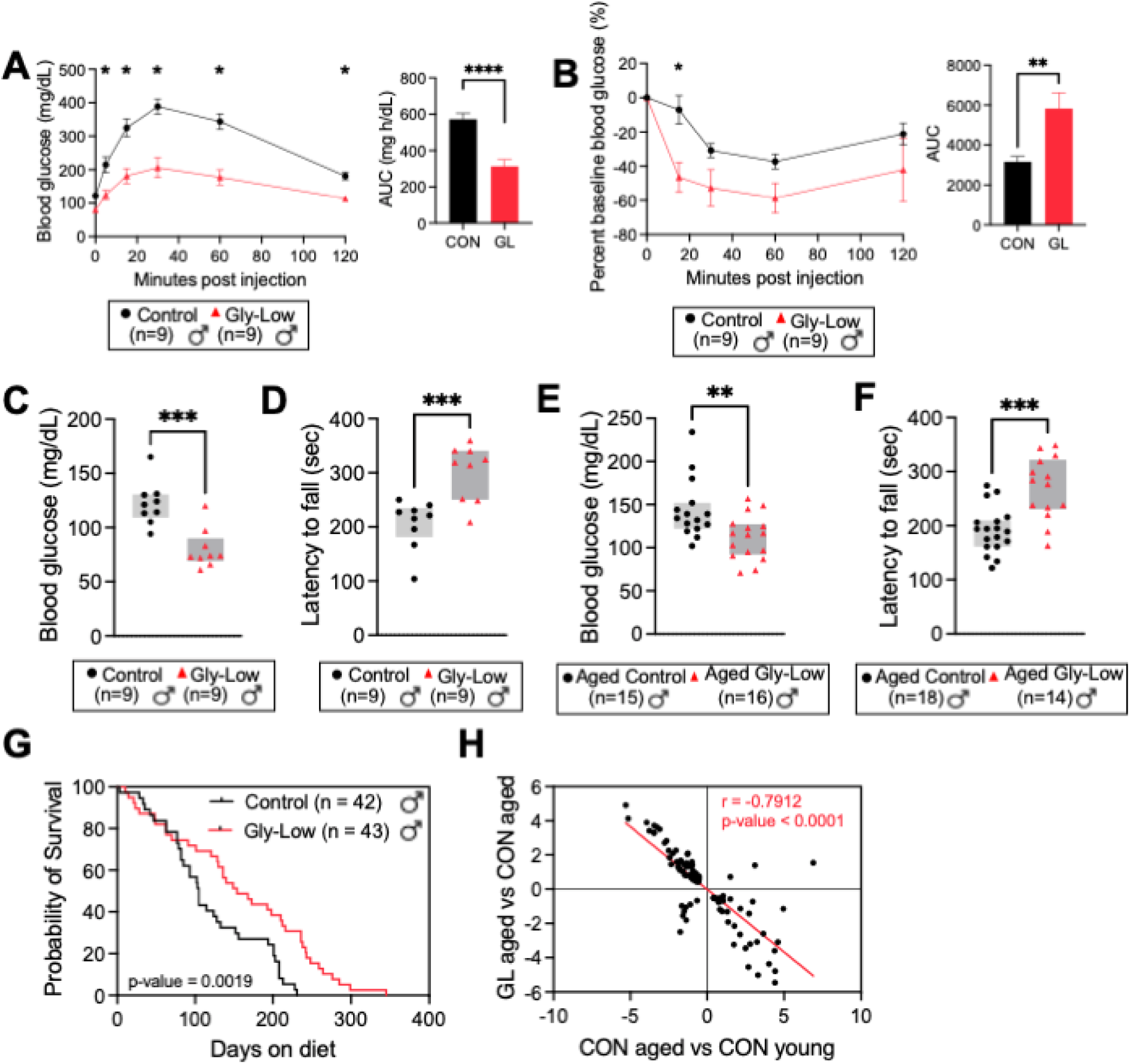
Impaired hypothalamic ghrelin signaling-induced mTOR activation with Gly-Low treatment, increases ribosomal translation, and shows potential as an anti-aging agent in middle-aged and old mice. A) Gly-Low treated mice had significantly reduced spikes in glucose levels during a glucose tolerance test (GTT) compared to control mice. AUC analysis (right). B) Glucose levels were decreased to a greater degree in mice treated with Gly-Low than control mice when administered a bolus of insulin during insulin tolerance testing (ITT). AUC analysis (right). Fasted blood glucose levels were reduced in middle-aged (12 month) (C) and aged (25 month) (E) wild-type male mice treated with Gly-Low compared to control mice. Rotarod performance was improved in middle-aged (12 month) (D) and aged (25 month) (F) Gly-Low treated mice compared to control mice. G) Kaplan-Meier curve showing lifespan extension as a late-life intervention (beginning at 24 months of age) in wildtype male Gly-Low treated compared to control mice. H) Scatterplot showing the log2 fold changes of genes significantly (p value < 0.05) altered in the hypothalamus of aged (25 month) Gly-Low treated vs aged (25 month) control, and aged (25 month) control vs young (3 month) control mice. The regression line is shown in red. The Pearson correlation coefficient, r, is shown in the top right quadrant. Significance: * p<0.05, ** p<0.005, **** p<0.00005. Statistical analyses performed by unpaired T-test.

### Gly-Low Rescues Aging-Associated Dysregulations in Glucose Homeostasis, Motor Coordination, and Extends Lifespan as a late-life treatment

Glycation stress increases with age accelerating numerous aging-associated conditions^57–59^. To test whether Gly-Low could influence age-related decline, we assessed glucose metabolism and motor function in control and Gly-Low treated middle-aged (12 months) and aged (20 months) wild-type male mice. Similar to improved glucose homeostasis observed in *Lepr^db^* mice, middle-aged male wild-type mice showed improved glucose metabolism during a glucose tolerance test. Following an injection of exogenous glucose, Gly-Low treated mice demonstrated a reduced peak in blood glucose levels relative to control mice (47.0% lower maximum blood glucose with a 45.4% lower area under the curve (AUC) value) (Figure 4A). Additionally, when these mice were later tested for their insulin response, Gly-Low treated mice responded more robustly. Gly-Low treated mice showed a 62.7% reduction in blood glucose relative to baseline levels, while control mice showed a 33.5% reduction (Figure 4B). Wild-type C57B6/J mice develop dysregulated glucose homeostasis with age and this is reflected in increased fasted and non-fasted blood glucose levels^60^. We found that Gly-Low reduced fasted blood glucose levels in both middle-aged and aged mice (Figure 4C and 4E). To test if Gly-Low was effective against age-related functional decline, we assessed motor function coordination in our middle-aged and aged mice through rotarod testing. The rotarod test is often used before behavioral or cognitive studies to identify sensorimotor impairments that may affect other assays^61^. Additionally, the rotarod test is commonly used to assess neuromuscular function, which declines with age^62,63^. We found that Gly-Low treated middle-aged and aged mice had improved performance on the rotarod test (Figure 4D and 4F). These findings indicate that Gly-Low prevented age-related decline in motor coordination and glucose homeostasis.

With their improved functional aging assays, we tested whether Gly-Low could influence lifespan as a late-life treatment. We began a cohort of male C57BL/6J mice with a Gly-Low diet at 24 months of age. Natural lifespan and deaths were recorded, and mice were only removed from the study based on stringent health parameters. Survival analysis demonstrated that control mice lived a median of 825 days (105 days after beginning dietary intervention) while Gly-Low treated mice lived a median of 888 days (173 days after beginning dietary intervention) (Figure 4G). This represented an 8.25% increase in overall median lifespan (p=0.0199, Mantel-Cox), and a 5.23% increase in maximum lifespan when treatment started at 24 months of age. This also translates to a difference in lifespan of 60.7% increase in time survived post-intervention. This extension in both median and maximum lifespan as a late-life intervention is not observed with caloric restriction (CR), suggesting that the extension is likely not driven solely by reduced food intake.^64,65^.

Hypothalamic aging has been shown to drive whole-body aging^66–70^. We assessed transcriptional differences between the hypothalamus of aged (25 months) Gly-Low treated and aged (25 months) control mice. From the differentially expressed genes found in this dataset, we performed linear regression against the genes identified when we compared aged control mice to young (3-month) control mice (Figure 4H). This regression shows a negative correlation (r = 0.8, p-value < 0.0001) between the two comparisons, suggesting that some changes in hypothalamic transcription seen in normal aging in the hypothalamus are reversed with a late-life intervention of Gly-Low.

## Discussion

Here, we report preclinical findings that support Gly-Low as a potential therapeutic tool for the treatment of obesity- and diabetes-associated pathologies. We selected the compounds that make Gly-Low based on their ability to protect against glycation stress. For that reason, we chose to initially test Gly-Low’s therapeutic potential in the well-studied, glycation-burdened *Lepr^db^* mouse model of hyperphagia and obesity^21^. As hypothesized, Gly-Low significantly reduced systemic levels of glycation stress as measured by reduced levels of the glycation precursor, methylglyoxal (MGO), and its arginine-adduct AGE, MG-H1. This study in mice and our previous studies in *C. elegans* support the use of glycation lowering compounds as a therapeutic strategy against diabetic pathologies^18,71^. MGO has long been suggested as a critical target for aging and age-related diseases^57–59^, but has been a challenge to target pharmacologically^71^. We demonstrate that Gly-Low reduces MGO and MG-H1 and that it likely does so by multiple mechanisms, including lowering production via glycolysis and increasing clearance by enhancing cellular detoxification pathways.

In addition to Gly-Low’s ability to reduce glycation stress, we found it also had appetite-suppressing effects resulting in reduced calorie intake. Given that *Lepr^db^* mice are glycation burdened and hyperphagic, we were not surprised to find that Gly-Low rescued many pathological phenotypes in this mouse model. However, Gly-Low also had health-promoting effects in young, middle-aged, and aged wild-type mice. Gly-Low’s therapeutic effect is likely a combination of its glycation-lowering capacity and calorie-reducing effect, which are intertwined. For example, we found that feeding worms MGH1 increases feeding^16^. Furthermore, feeding mice MGO-modified bovine serum albumin induced insulin resistance, increased body weight, and shortened lifespan^72^. A clinical trial in which people were given glyoxalase activating compounds, which detoxify MGO, found that these compounds reduced weight and insulin resistance ^27^. These findings suggest that Gly-Low’s health-promoting effects are likely due to a complex interaction between its ability to directly reduce glycation stress and directly or indirectly reduce food consumption. The mechanism by which Gly-Low affects feeding behavior potentially involves multiple mechanisms and includes suppression of the appetite-stimulating ghrelin pathway and activation of the appetite-satiating mTOR pathway within the hypothalamus.

Caloric restriction (CR) is one of the most potent and widely conserved interventions for increasing healthspan and lifespan across species^73^. Some of Gly-Low’s health-promoting effects are likely due to its calorie-reducing effect. However, in this study, we demonstrated that Gly-Low may separate itself from conventional CR. One important distinction between CR and Gly-Low is Gly-Low’s ability to extend lifespan by ∼60% even when administered as a late-life treatment. The late-life benefits of CR are generally titrated by the age at which CR is initiated^64,65^. A report by Lipman in 1995 demonstrates no change in median lifespan of rats fed 33% less than controls (33% CR) when treated beginning at 18 months of age^45^. Interestingly, we demonstrate in our study that Gly-Low treatment at 24 months of age resulted in an increase in both median and maximum lifespan, with mice voluntarily eating 13.4-29.6% less than control mice. While our study cannot be directly compared to traditional CR studies due to the differences in caloric restriction rates, it will be worth investigating further in the future. We hypothesize that Gly-Low is promoting metabolic health with anti-aging effects in part through altered hypothalamic signaling, which in parallel, influences feeding behavior and whole body aging phenotypes^66–69^. Hypothalamic transcript analysis from Gly-Low treated aged mice indicates that many genes whose expression changes with age are reversed when old mice are treated with Gly-Low (Figure 4H).

Caloric restriction (CR) has been shown to improve metabolic health and slow aging and age-related diseases in multiple species^74^. However, long-term CR is not sustainable in humans^75^. Lowering glycation stress with therapies such as Gly-Low holds promise to induce a voluntary reduction in food consumption, independently ameliorate tissue pathologies, and slow aging-related outcomes that result from an increase in glycation stress due to methylglyoxal and related precursors.

## Author contributions

PK, SK, JG, JCN, LE involved in conceptualization and supervision. LW, KK, PK, LE, JCN drafted the manuscript. LW, KK involved in analysis and visualization. LW, KK, JR, NB, MV, MMS, DF, PS, JB, DS, LEN involved in experimental procedure, live animal experimentation. SJC involved in blinded pathological assessment. JG and DF involved in experimental quantification and analysis of α-dicarbonyls. All authors have read and agreed to the final version of the manuscript.

## Acknowledgements

We thank A. Bartke and D. Medina for discussion and experimental procedure development, B. Schilling and S. Melov for useful discussion, A. Lopez-Ramirez for brain dissections and contribution to the work, and C. Patterson for contribution to cohort husbandry and experimentation. We thank the Kapahi Lab at Buck Institute for helpful discussion and critique, and we thank the employees of the Phenotyping Core and Morphology Core at the Buck Institute.

## Data availability

The data supporting this study’s findings are available from the corresponding author upon request.

## Inclusion and diversity

We support inclusive, diverse, and equitable conduct of research.

## Methods

### Mice

Studies included the use of male C57BL/6J (Jackson Laboratories #000664) control mice, male leptin receptor deficient mice (Jackson Laboratories #000642, homozygous for *Lepr^db^*, wild type for Dock7^m^) aged between 8 and 12 weeks, as well as 19-month-old male C57BL/6J mice (National Institutes of Aging, Bethesda, MD). All mice were communally housed and age-matched with *ad libitum* access to water and diet in a pathogen and temperature-controlled room with a 12h light-dark cycle beginning at 06:00 AM. All procedures were conducted per NIH Guidelines for Care and Use of Animals and were approved by the Institutional Animal Care and Use Committees at Buck Institute for Research on Aging and the University of California San Francisco.

### Experimental Diets

Mice were fed either a standard low-fat chow diet (21% fat (kcal), 60% carbohydrate (kcal) Envigo: TD.200743), a standard high fat chow diet (60% fat (kcal), 21% carbohydrate (kcal), Envigo: TD.200299), a standard low-fat chow diet supplemented with our Gly-Low compound cocktail (21% fat (kcal), 60% carbohydrate (kcal) Envigo: TD.200742), or a standard high fat chow diet supplemented with our Gly-Low compound cocktail (60% fat (kcal), 21% carbohydrate (kcal).

A combination of supplemental grade compounds, safe to be consumed in set dosages, were prepared and incorporated into a modified pre-irradiated standard AIN-93G mouse chow diet from Envigo. The cocktail consists of alpha lipoic acid (20.19%), nicotinamide (57.68%), thiamine hydrochloride (4.04%), piperine (1.73%), and pyridoxamine dihydrochloride (16.36%), and is supplemented in the diet to achieve a daily consumption rate in mg/kg of body weight/day. For the 1X (full-dose) diet, these percentages translate to 3 g/kg alpha lipoic acid, 8.57 g/kg nicotinamide, 0.6 g/kg thiamine hydrochloride, 0.26 g/kg piperine, and 2.43 g/kg pyridoxamine dihydrochloride. Noted vocabulary where 1X is equal to a full dose, 0.5X is equal to a half dose, and 0.25X is equal to a quarter dose of the compound cocktail.

### Lifespan

19-month-old male C57BL/6J mice received from National Institutes of Aging (Bethesda, MD) and maintained on vivarium chow until they reached 24 months of age. At 24 months of age, mice were randomly assigned to begin either a control chow diet or Gly-Low diet. Health checks were performed 5 times a week and mice reached either a humane endpoint or died of natural causes.

### Measuring Food Intake and Energy Metabolism

Food consumption and body weight were measured once weekly. Chow weight of communally caged mice was recorded once weekly and individual intake was measured as change in weight over the week divided by the number of mice per cage.

Metabolic parameters in mice were assessed using a Promethion Metabolic Caging System housed in the Buck Institute Mouse Phenotyping Core. Mice were singly housed and received water and food *ad libitum*. Cages were maintained at 20°-22° C under a 12h light-dark cycle beginning at 06:00 AM, and mice were acclimated to single housing 24h before being studied. The cages continuously weighed food for each mouse, and daily intake was measured as a change in food weight over 24h periods. Metabolic data were collected by the respiration rates of each mouse and were normalized to individual mouse body mass. X, Y, and Z beam breaks quantified total activity and steps taken during the metabolic cage run.

### Body Composition Quantification

Echo MRI and DXA scans were used to analyze body composition in anesthetized (isoflurane) immobilized mice. Water weight and bone mass were excluded from body weight to quantify lean and fast mass.

### Glucose Testing

Random (non-fasted) and fasted blood glucose levels were determined for each mouse after 8 weeks and 16 weeks of treatment. Non-fasted blood glucose levels were assessed between 08:00 AM and 10:00 AM by a collection of blood by tail nick and the use of a handheld glucometer (AccuCheck). Fasted blood glucose was determined after a 16h fast between 8:00 AM and 10:00 AM by a collection of blood by tail nick and the use of a handheld glucometer (AccuCheck). Handheld Accucheck glucometer had a maximum read capacity of 600 mg/dL. Therefore, any values above max read capacity were listed as 600 mg/dL.

### Glucose and Insulin Tolerance Testing

Mice underwent GTT testing at 12 months of age, following 8 months of Gly-Low treatment. Food was removed from control-fed and Gly-Low treated mice for 14 hr before testing of glucose tolerance from 6:00 PM to 8:00 AM. Mice received a single IP injection of D-glucose (2 g/kg), followed by a single tail nick to collect blood for blood glucose monitoring by handheld glucometer (AccuCheck). Total glucose AUC was measured by GraphPad Analysis.

Mice underwent ITT testing at 12 months of age, following 8 months of Gly-Low treatment. Food was removed from control-fed and Gly-Low treated mice 4 hr before testing of insulin tolerance from 8:00 AM to 12:00 PM. Mice received IP injections of insulin (0.75U/kg), followed by a single tail nick to collect blood glucose monitoring by handheld glucometer (AccuCheck). Mice that had a blood glucose reading <40 mg/dL were immediately removed from the study and injected with 100 uL of 2 g/kg D-glucose. Total glucose AUC was measured by GraphPad Analysis. Handheld Accucheck glucometer had a maximum read capacity of 600 mg/dL. Therefore, any values above max read capacity were listed as 600 mg/dL.

### Leptin Injections

Mice underwent leptin sensitivity testing at 4 months of age, following 1 month of Gly-Low treatment. Food was removed from control-fed and Gly-Low treated mice 16 hr before testing of leptin sensitivity from 7:00 AM to 7:00 PM. Mice received a single IP injection of either saline vehicle or leptin (1 g/kg). Food consumption rates were collected by metabolic cage food measurements. Mice were allowed four days of rest before undergoing another single IP injection of either saline vehicle or leptin (1 g/kg) to serve as their control. Total food consumption was measured by GraphPad Analysis.

### Leptin Injection and pSTAT3 Signaling

Mice underwent histological leptin sensitivity testing at 4 months of age, following 1 month of Gly-Low treatment. Food was removed from control-fed and Gly-Low treated mice 16 hr before testing of leptin sensitivity from 7:00 AM to 7:00 PM. Mice received a single IP injection of either saline vehicle or leptin (1 g/kg). Within 30 min of receiving the injection, mice were euthanized by CO_2_ asphyxiation and cervical dislocation. Dissections were performed and mouse brains were removed and washed with PBS before being postfixed in 4% PFA overnight with agitation at 4C. Afterward, a brain matrix (BrainTree) was used to isolate sections containing the hypothalamus. This section was immediately embedded in OCT, frozen on dry ice, and stored at −80C. Next, 4-micron sections were cut on a cryostat, blocked for 1 hr with 5% BSA containing 0.1% triton X-100, and incubated with pSTAT3 (1:200, Cell Signaling). Adequate secondary antibody was used for the HRP-diaminobenzidine reaction. The HRP-diaminobenzidine reaction was performed using the ABC Kit (Vector Laboratories), using biotin-labeled goat anti-rabbit IgG. Images were acquired using a Zeiss AxioImager brightfield microscope.

### Ghrelin Responsiveness

Ghrelin peptides (Catalog #031-30)(Phoenix Pharmaceuticals, Inc.). Mice were injected with reconstituted ghrelin (0.1 mg/kg) by subcutaneous injection. Following injection, mice were singly housed, and individual food consumption was recorded over 90 minutes post-injection.

#### Hormone Quantification

Blood samples were collected by cardiac puncture when mice were euthanized for dissection. Blood samples were collected in heparin lined tubes and left on ice for 30 min. Afterward, samples were centrifuged for 15 min at 2200 g to isolate plasma. Ghrelin, LEAP2, and IGF-1 were measured in plasma by ELISA kits according to manufacturer’s instructions.

LEAP2 ELISA (Catalog #075-40) (Phoenix Pharmaceuticals, Inc.)

Ghrelin ELISA (EZRGRA-90K)(Sigma Aldrich)

IGF-1 ELISA (Catalog #80574) (Crystal Chem)

#### Hypothalamic RNA Sequencing

Hypothalamic transcripts were analyzed from male C57BL/6J mice at three ages under different treatment paradigms: 1) young (4-month-old) male C57BL/6J mice fed a control or glycation-lowering (Gly-Low) diet for 1 week, 2) aged (25-month-old) male C57BL/6J mice fed a control or glycation-lowering (Gly-Low) diet for 5 months and 3) young (3-month-old) male C57BL/6J mice fed a control diet for 1 week. Aged 19 month-old male C57BL/6J mice were ordered from the National Institutes of Aging (Bethesda, MD). Young mice were acquired from Jackson laboratories (#000664). Mice were fed vivarium chow (Envigo Teklad 2018) before starting either a control diet or Gly-Low diet. Mice were sacrificed via CO_2_ asphyxiation followed by cervical dislocation. The brain was rapidly dissected and the hypothalamus was removed with tweezers, flash-frozen, and stored at −80°C. RNA was isolated using Zymo research quick RNA miniprep kit (cat # 11-328) according to the manufacturer’s recommendations. Isolated RNA was sent for library preparation and sequencing by Novogene Corporation Inc. where RNA was poly-A selected using poly-T oligo-attached magnetic beads, fragmented, reverse transcribed using random hexamer primers followed by second strand cDNA synthesis using dTTP for non-directional library preparation. Samples underwent end repair, A-tailing, adapter ligated, size selected, amplified, and purified. Illumina libraries were quantified using Qubit and qPCR and analyzed for size distribution using a bioanalyzer. Libraries were pooled and sequenced on an Illumina Novoseq 6000 to acquire paired-end 150 bp reads. Data quality was assessed and adaptor reads and low quality reads were removed. Reads that passed the quality filtering process were mapped paired-end to the reference genome (GRCm38) using Hisat2 v2.0.5. featureCounts v1.5.0-p3 was used to count reads that mapped to each gene. Differential expression analysis was performed using DESEq2 (1.20.0). Where indicated, bootstrapping was performed using R (R version 4.1.2) program ‘boot’ (1.3-28.1). To determine the expected mean and standard deviation, n=i log2 fold changes were randomly selected 1000 times, in which i is the number of genes in the gene set.

#### Metabolomics

Samples were prepared and analyzed by Norwest Metabolomics Research Center according to the following:

**LC-MS Conditions:** *Acquisition Mode MS1*; threshold count: 100; *m/z* 60-1000; Gas temp 325°C; Drying Gas 10L/min; Nebulizer 35 psi; TOF Fragmentator 120 V; Skimmer 65 V; 4 spectra/s; 250 ms/spectrum; *Acquisition Mode MS2:* range *m/z* 20-1000; 4 spectra/s; 250 ms/spectrum; CE 20 eV; 5 precursor per cycle, MS2 threshold 200 counts; active exclusion after 3 spectra; static exclusion rang m/z 60-100; Abundance dependent accumulation target 50000 counts/spectrum; exclusion list from blank injection enabled; reference mass correction enabled.

**LC System:** Agilent 1260 as the Mobile Phase (pump 2) and reference solution (pump 1)

**MS System:** Agilent 1200 SL LC-6520 Quadrupole-Time of Flight (Q-TOF) MS

**LC column**: WATERS XBridge BEH Amide (15 cm x 2.1 mm; 2.5 µm)

**Buffer A:** 10 mM ammonium acetate and 0.2% acetic acid in 95% H_2_O + 3% ACN + 2% MeOH;

**Buffer B:** 10 mM ammonium acetate and 0.2% acetic acid in 5% H_2_O + 93% ACN + 2% MeOH;

**Injection**: 5 μL (+)-ESI and 10 μL (-)-ESI; **Wash:** 95%ACN+5%H_2_O for 10 s; **flow rate** (mL/min): 0.3

**LC Column Chamber Temp:** 40° C; **ESI mode:** (+/-)

**Worklist:** each sample was injected in both positive and negative ESI mode (labeled _POS and _NEG). Blank of sample preparation labeled as blank_Prep. Plasma and tissue samples were combined to make QC samples from plasma (QCp) and tissue (QCt).

### Gradient operation (Separation)

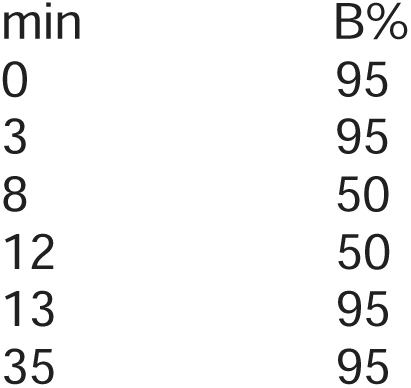

#### Plasma sample preparation

Samples were thawed at 4°C, vortexed 10 seconds. 28uL plasma plus 22uL water was transferred to a 2 mL Eppendorf vial. 250uL Methanol was added then the sample was vortexed 10 seconds. Samples were incubated at −20°C for 20 minutes and centrifuged at 14000 rpm at 4°C for 15 minutes. 150uL supernatant was transferred to a new 2 mL Eppendorf vial and dried completely using a Vacufuget at 30°C for about 1.5 hours. Samples were reconstituted with 250uL HILIC solvent, vortexed 10 seconds then centrifuged for 5 min at 14000 rpm at 4°C. 250uL supernatant was transferred into LC vials for MS analysis.

#### Data processing

Data processing was performed using Progenesis Qi software v. 2.2.5826.42898 (Nonlinear Dynamics; Newcastle; UK). Peak alignment was carried out taking samples as reference. Peak-peaking was performed using sensitivity and chromatographic peak width at 1 (3 for (+)-ESI) and 0.01min; respectively. The retention time limit was set 1.0 – 12.0 min. Possible adduct ions were defined as follows: [M+H]^+^; [M+Na]^+^; [M+NH_4_]^+^; [M+K]^+^; M^+●^; [M+ACN+H]^+^; in (+) ESI mode; and [M–H]^−^; [M+HCOO–H]^−^; [M+Cl]^−^; and [2M+CH3COO–H]^−^ in (-) ESI mode. The adduct ions were grouped into mass features through peak deconvolution. Putative peak annotation was performed by searching metabolites from MONA Database using accurate *m/z*measurements from the full scan data and MS2 spectra. We set the *m/z* tolerance of 20 ppm (MS1) and 30 ppm (MS2).

#### Liver Histology

Dissections were performed and mouse livers were removed and washed with PBS before being postfixed in 4% PFA overnight with agitation at 4C. Afterwards, livers were moved to 70% EtOH before being paraffin embedded and sectioned. The liver was sectioned by microtome in a coronal orientation at a thickness of 4 microns. H&E staining and trichrome staining were used for the identification and quantification of large lipid vacuoles.

#### Western Blotting

Proteins were extracted from hypothalamic samples in TPER buffer (Thermo Fisher) containing a protease inhibitor cocktail (Sigma). Protein extracts were denatured at 70°C for 15 min prior to running on a 5-12% Bis-Tris gel. The transfer was completed using iBlot (Thermo Fisher) and blocked in 5% BSA for 1 hr. Primary antibodies were incubated overnight at 4°C with agitation. Adequate HRP-conjugated antibodies were incubated at room temperature for 1 hr prior to imaging.

Akt (Cell Signaling)(40D4) 1:1000

pAKT Ser473 (Cell Signaling)(D9E) 1:1000

S6: (Cell Signaling) (5G10) (2217) 1:1000

pS6 Ser 240/244: (Cell Signaling) (D68F8) 1:1000

AMPK: (Cell Signaling) (CST-4181) 1:1000

pAMPK: (Cell Signaling) (CST-2531)

mTOR: (Cell Signaling) (7C10): 1:1000

Actin (Cell Signaling) (13E5) 1:1000

#### Mass Spectrometry MGO Quantification

##### Quantification of Dicarbonyls

200 µL of 80:20 MeOH:ddH_2_O (−80°C) containing 50 pmol ^13^C_3_-MGO was added to 10 µL of serum and extracted at −80°C overnight. Insoluble protein was removed via centrifugation at 14,000 x g for 10 min at 4°C. Supernatants were derivatized with 10 µL of 10 mM o-phenylenediamine for 2 h with end-over-end rotation protected from light.^76^. Derivatized samples were centrifuged at 14,000 x *g* for 10 min, and the supernatant was chromatographed using a Shimadzu LC system equipped with a 150 x 2mm, 3μm particle diameter Luna C_18_ column (Phenomenex, Torrance, CA) at a flow rate of 0.450 mL/min. Buffer A (0.1% formic acid in H_2_O) was held at 90% for 0.25 min, then a linear gradient to 98% solvent B (0.1% formic acid in acetonitrile) was applied over 4 min. The column was held at 98% B for 1.5 min, washed at 90% A for 0.5 min, and equilibrated to 99% A for 2 min. Multiple reaction monitoring (MRM) was conducted in positive ion mode using an AB SCIEX 4500 QTRAP with the following transitions: *m/z* 145.1→77.1 (MGO); *m/z* 235.0→157.0 (3-DG); m/z 131.0→77.0 (GO); m/z 161.0→77.0 (HPA); m/z 148.1→77.1 (^13^C_3_-MGO, internal standard).

##### Quantitation of PTMs (QuarkMod)

Protein pellets from the dicarbonyl quantifications (above) were resuspended in 65 µL of 50 mM NH_4_HCO_3_, pH 8.0. Samples were spiked with 10 μL of a master mix containing internal standards (see table). Proteins were digested by adding 5 μL of sequencing grade trypsin (0.1 mg/mL) (Promega) for three h at 37 °C. Trypsin was denatured by boiling at 95 °C for 10 minutes, and the samples were cooled to room temperature. Aminopeptidase M (Millipore, 15 µg in 10 µL) was added, and samples were incubated overnight at 37°C. Aminopeptidase was denatured via heating at 95 °C for 10 min, and samples were again cooled to room temperature. 15 µL of heptafluorobutyric acid (1:1 in H_2_O) was added to each sample, and debris was removed via centrifugation at 14,000 × *g* for 10 min. Clarified supernatants were chromatographed using a Shimadzu LC system equipped with a 150 x 2.1mm, 3.5 mm particle diameter Eclipse XDB-C8 column (Agilent, Santa Clara, CA) at a flow rate of 0.4500 mL/min. Mobile phase A: 10 mM HFBA in water; mobile phase B: 10 mM HFBA in ACN. The following gradient was used: 2 min, 1% B; 6 min, 50% B; 6.5 min, 95% B; 9 min, 95% B; 9.5 min, 1% B. The column was equilibrated for 3 min at 5% B. MRM was conducted in positive mode using an AB SCIEX 4500 QTRAP. The MRM detection window was 50 sec with a target scan time of 0.75 sec. The following parameters were used for detection and as previously described^77,78^:

| Species | Q1 (*m/z*) | Q3 (*m/z*) | CE (V) |
| --- | --- | --- | --- |
| Lys | 147.1 | 84.1 | 29 |
| $^{13}\text{C}_6^{15}\text{N}_2$ Lys | 155.1 | 90.1 | 29 |
| Arg | 175.1 | 70.1 | 47 |
| $^{13}\text{C}_6^{15}\text{N}_4$ Arg | 185.1 | 75.1 | 47 |
| Leu | 132.1 | 86.1 | 17 |
| $^{13}\text{C}_6^{15}\text{N}$ Leu | 139.1 | 93.1 | 17 |
| MG-H1 | 229.2 | 70.1 | 53 |
| $^{13}\text{C}$-MG-H1 | 230.2 | 70.1 | 53 |
| CEA | 247.2 | 70.1 | 55 |
| $^{13}\text{C}$-CEA | 248.2 | 70.1 | 55 |
| CEL | 219.2 | 84.1 | 41 |
| CEL-d<sub>4</sub> | 223.2 | 88.1 | 41 |
| CML | 205.0 | 84.1 | 38 |
| CML-d<sub>4</sub> | 209.0 | 88.1 | 38 |

##### Quantification of free MGH1

10 µL of serum was added to 200 µL of 80:20 MeOH:ddH_2_O (−80°C) containing ten pmol ^13^C-MG-H1 and extracted at −80°C overnight. Insoluble protein was removed via centrifugation at 14,000 x g for 10 min at 4°C, and supernatants were transferred to a new tube. 15 µL of heptafluorobutyric acid (1:1 in H_2_O) was added to each sample, and debris was removed via centrifugation at 14,000 × *g* for 10 min. Samples were analyzed as described above (QuARKMod).

#### Quantification and statistical analysis

Statistical details of experiments can be found in the figure legends. As the figure legends indicate, all data are expressed as mean ± SEM. Statistical tests were selected based on appropriate assumptions with respect to data distribution and variance characteristics. For normally distributed data, statistical significance was determined using an unpaired T-test. Statistical significance was determined using a one-sample T-test for data normalized to the vehicle control group. All statistical analyses were performed using GraphPad Prism. Significant differences are indicated: ^∗^ *p* ≤ 0.05, ^∗∗^ *p* ≤ 0.005, ^∗∗∗^ *p* ≤ 0.0005, ^∗∗∗∗^ *p* < 0.0001.

