## Supplementary for "Glycation-lowering compounds inhibit ghrelin signaling to reduce food intake, lower insulin resistance, and extend lifespan"

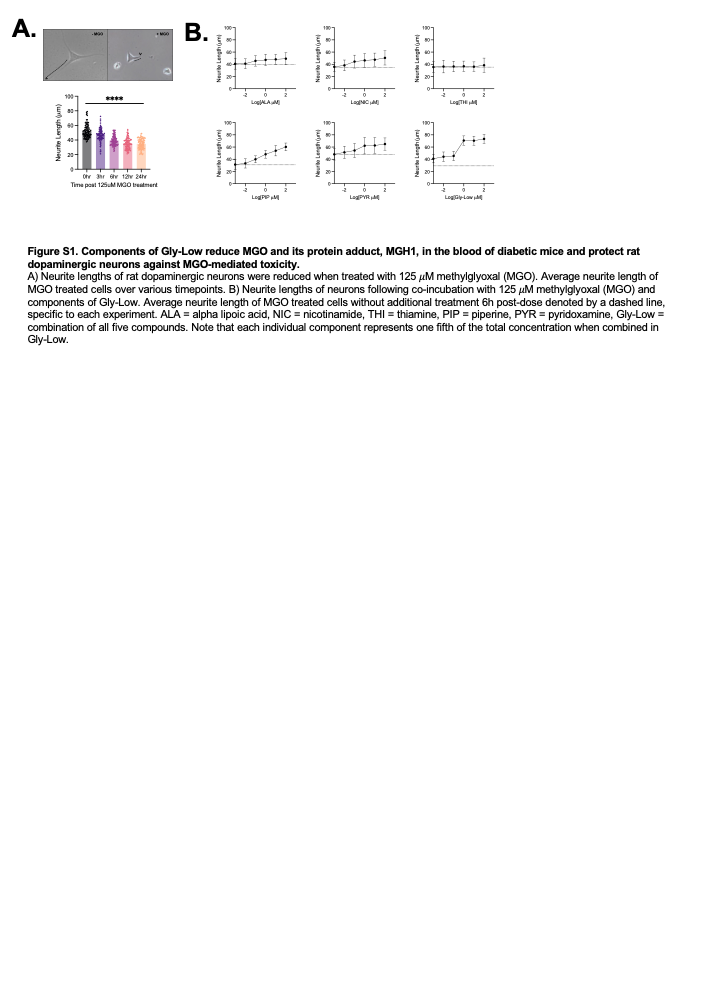

**Figure S1. Components of Gly-Low reduce MGO and its protein adduct, MGH1, in the blood of diabetic mice and protect rat dopaminergic neurons against MGO-mediated toxicity.**

A) Neurite lengths of rat dopaminergic neurons were reduced when treated with 125 𝜇M methylglyoxal (MGO). Average neurite length of MGO treated cells over various timepoints. B) Neurite lengths of neurons following co-incubation with 125 𝜇M methylglyoxal (MGO) and components of Gly-Low. Average neurite length of MGO treated cells without additional treatment 6h post-dose denoted by a dashed line, specific to each experiment. ALA = alpha lipoic acid, NIC = nicotinamide, THI = thiamine, PIP = piperine, PYR = pyridoxamine, Gly-Low = combination of all five compounds. Note that each individual component represents one fifth of the total concentration when combined in Gly-Low.

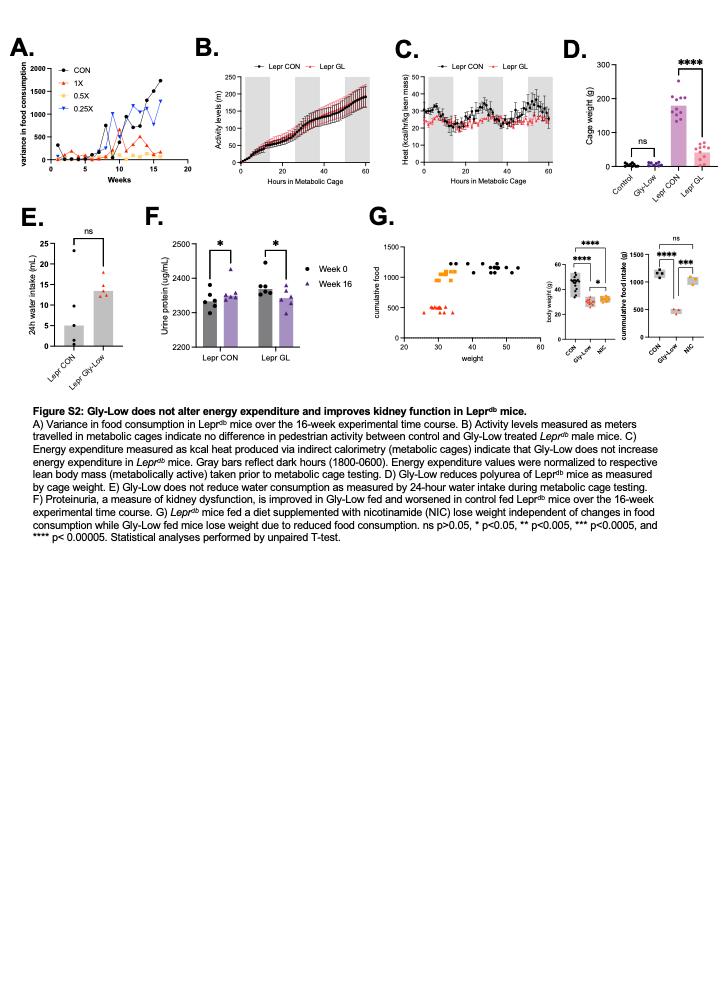

**Figure S2: Gly-Low does not alter energy expenditure and improves kidney function in Lepr^db^ mice.**

A) Variance in food consumption in Lepr^db^ mice over the 16-week experimental time course. B) Activity levels measured as meters travelled in metabolic cages indicate no difference in pedestrian activity between control and Gly-Low treated *Lepr^db^* male mice. C) Energy expenditure measured as kcal heat produced via indirect calorimetry (metabolic cages) indicate that Gly-Low does not increase energy expenditure in *Lepr^db^* mice. Gray bars reflect dark hours (1800-0600). Energy expenditure values were normalized to respective lean body mass (metabolically active) taken prior to metabolic cage testing. D) Gly-Low reduces polyurea of Lepr^db^ mice as measured by cage weight. E) Gly-Low does not reduce water consumption as measured by 24-hour water intake during metabolic cage testing. F) Proteinuria, a measure of kidney dysfunction, is improved in Gly-Low fed and worsened in control fed Lepr^db^ mice over the 16-week experimental time course. G) *Lepr^db^* mice fed a diet supplemented with nicotinamide (NIC) lose weight independent of changes in food consumption while Gly-Low fed mice lose weight due to reduced food consumption. ns p>0.05, * p<0.05, ** p<0.005, *** p<0.0005, and **** p< 0.00005. Statistical analyses performed by unpaired T-test.

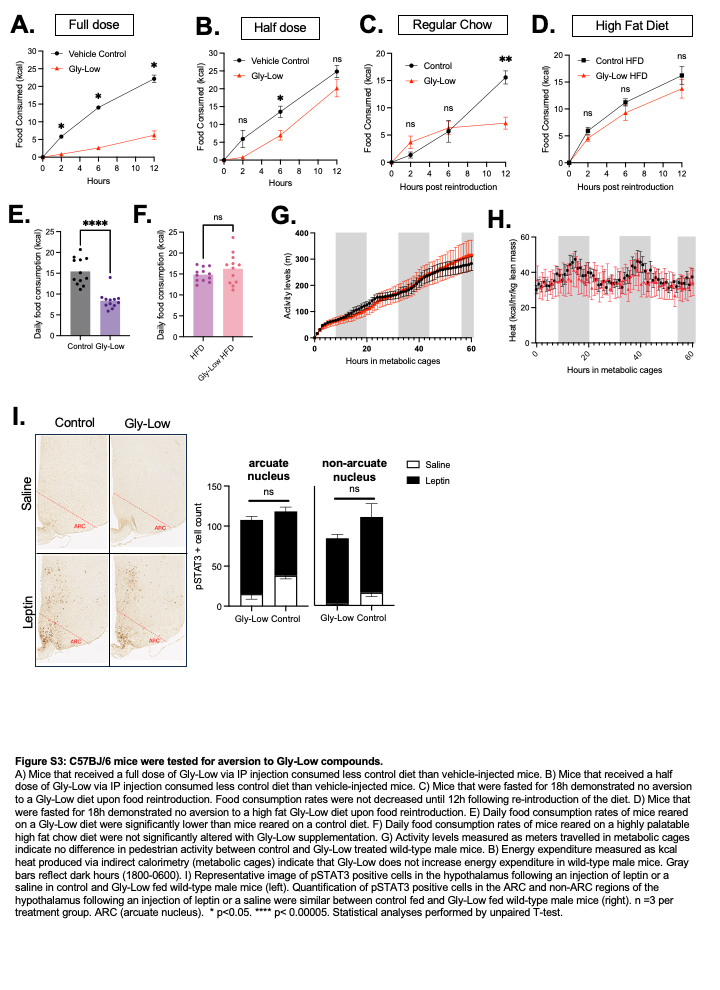

**Figure S3: C57BJ/6 mice were tested for aversion to Gly-Low compounds.**

A) Mice that received a full dose of Gly-Low via IP injection consumed less control diet than vehicle-injected mice. B) Mice that received a half dose of Gly-Low via IP injection consumed less control diet than vehicle-injected mice. C) Mice that were fasted for 18h demonstrated no aversion to a Gly-Low diet upon food reintroduction. Food consumption rates were not decreased until 12h following re-introduction of the diet. D) Mice that were fasted for 18h demonstrated no aversion to a high fat Gly-Low diet upon food reintroduction. E) Daily food consumption rates of mice reared on a Gly-Low diet were significantly lower than mice reared on a control diet. F) Daily food consumption rates of mice reared on a highly palatable high fat chow diet were not significantly altered with Gly-Low supplementation. G) Activity levels measured as meters travelled in metabolic cages indicate no difference in pedestrian activity between control and Gly-Low treated wild-type male mice. B) Energy expenditure measured as kcal heat produced via indirect calorimetry (metabolic cages) indicate that Gly-Low does not increase energy expenditure in wild-type male mice. Gray bars reflect dark hours (1800-0600). I) Representative image of pSTAT3 positive cells in the hypothalamus following an injection of leptin or a saline in control and Gly-Low fed wild-type male mice (left). Quantification of pSTAT3 positive cells in the ARC and non-ARC regions of the hypothalamus following an injection of leptin or a saline were similar between control fed and Gly-Low fed wild-type male mice (right). n =3 per treatment group. ARC (arcuate nucleus). * p<0.05. **** p< 0.00005. Statistical analyses performed by unpaired T-test.

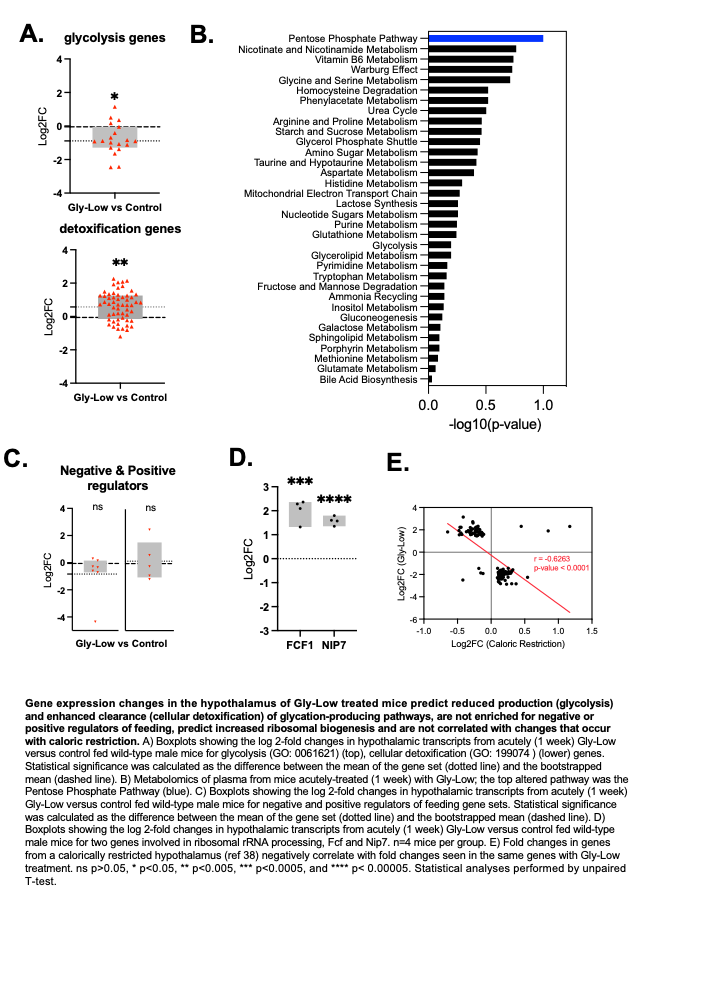

**Gene expression changes in the hypothalamus of Gly-Low treated mice predict reduced production (glycolysis) and enhanced clearance (cellular detoxification) of glycation-producing pathways, are not enriched for negative or positive regulators of feeding, predict increased ribosomal biogenesis and are not correlated with changes that occur with caloric restriction.** A) Boxplots showing the log 2-fold changes in hypothalamic transcripts from acutely (1 week) Gly-Low versus control fed wild-type male mice for glycolysis (GO: 0061621) (top), cellular detoxification (GO: 199074 ) (lower) genes. Statistical significance was calculated as the difference between the mean of the gene set (dotted line) and the bootstrapped mean (dashed line). B) Metabolomics of plasma from mice acutely-treated (1 week) with Gly-Low; the top altered pathway was the Pentose Phosphate Pathway (blue). C) Boxplots showing the log 2-fold changes in hypothalamic transcripts from acutely (1 week) Gly-Low versus control fed wild-type male mice for negative and positive regulators of feeding gene sets. Statistical significance was calculated as the difference between the mean of the gene set (dotted line) and the bootstrapped mean (dashed line). D) Boxplots showing the log 2-fold changes in hypothalamic transcripts from acutely (1 week) Gly-Low versus control fed wild-type male mice for two genes involved in ribosomal rRNA processing, Fcf and Nip7. n=4 mice per group. E) Fold changes in genes from a calorically restricted hypothalamus (ref 38) negatively correlate with fold changes seen in the same genes with Gly-Low treatment. ns p>0.05, * p<0.05, ** p<0.005, *** p<0.0005, and **** p< 0.00005. Statistical analyses performed by unpaired T-test.
